# Photoacoustic molecular rulers based on DNA nanostructures

**DOI:** 10.1101/125583

**Authors:** James Joseph, Philipp Koehler, Tim J. Zuehlsdorff, Daniel J. Cole, Kevin N. Baumann, Judith Weber, Sarah E. Bohndiek, Silvia Hernández-Ainsa

**Author notes:** Corresponding Authors: Silvia Hernández-Ainsa, Sarah E. Bohndiek.

## Abstract

Molecular rulers that rely on the Förster resonance energy transfer (FRET) mechanism are widely used to investigate dynamic molecular processes that occur on the nanometer scale. However, the capabilities of these fluorescence molecular rulers are fundamentally limited to shallow imaging depths by light scattering in biological samples. Photoacoustic tomography (PAT) has recently emerged as a high resolution modality for *in vivo* imaging, coupling optical excitation with ultrasound detection. In this paper, we report the capability of PAT to probe distance-dependent FRET at centimeter depths. Using DNA nanotechnology we created several nanostructures with precisely positioned fluorophore-quencher pairs over a range of nanoscale separation distances. PAT of the DNA nanostructures showed distance-dependent photoacoustic signal generation and experimentally demonstrated the ability of PAT to reveal the FRET process deep within tissue mimicking phantoms. Further, we experimentally validated these DNA nanostructures as providing a novel and biocompatible strategy to augment the intrinsic photoacoustic signal generation capabilities of small molecule fluorescent dyes.

---

Nanoscale assessment of distance-dependent fluorescence quenching has been widely utilized in nanotechnology and biomedicine to investigate dynamic molecular processes that occur on the nanometer scale.^1, 2^ These fluorescence molecular rulers are fundamentally limited by light scattering in biological samples, which severely restricts the penetration depth for imaging due to the short (<100 μm) mean free path of photons in biological media.^3^ As a result, the application of fluorescence-based molecular rulers is typically limited to the study of cells *in vitro*^4^, ^*^5^*^ or to superficial applications with intravital microscopy *in vivo*.^6^ Although intravital imaging based on FRET^7^ mechanisms are being extensively used in various domains of cell biology^6, 8, 9^ and drug discovery^8, 10, 11^ they are challenged with limited penetration depth, low signal to noise ratio and photobleaching. Hence, there exists an unmet need for methodologies to probe dynamic molecular interactions that can reveal cellular responses at depth in intact living subjects.

Photoacoustic (PA) tomography (PAT) is emerging as an *in vivo* preclinical imaging tool that can overcome the traditional depth limitations of all-optical imaging, providing images with a resolution of ∼100 μm at depths of up to 3 cm.^12^ PAT is a hybrid modality based on the absorption of pulsed light in tissue, which generates a transient thermoelastic expansion and produces an acoustic wave that can be detected by ultrasound transducers at the tissue surface. PAT requires that the decay of the optical excitation occurs via non-radiative processes to provide thermalization of the absorbed energy.^13^ Fluorescence quenching is one such non-radiative decay that leads to heat dissipation into the surrounding medium.^14^ We therefore hypothesized that the presence of fluorescence quenching would translate into a corresponding enhancement of the PA signal, which would therefore enable distance-dependent fluorescence quenching to be measured at depth in living subjects.

DNA nanotechnology is well established as a tool that enables the construction of well-defined nanostructures with a range of structural and molecular functionalities.^15, 16^ Due to the accurate specificity of base-pair interactions, it is possible to obtain self-assembled DNA nanoplatforms that allow accurate positioning of various moieties with sub-nanometer precision.^17, 18^ The decoration of these DNA constructs with fluorophores^19-23^ has already resulted in several studies of molecular interactions based on FRET pairs using dye-dye or dye-quencher combinations. These pairs have been used extensively in DNA nanostructures as reporters for different purposes, including molecular probes,^24^ single-molecule studies,^25-28^ DNA machines^29, 30^ and DNA walkers.^31^

Here, we report a systematic study of PAT molecular rulers using DNA nanostructures to precisely tune the distance between a fluorophore and quencher pair suitable for *in vivo* imaging in the near-infrared (NIR) optical window.^32^ We assess the absorbance, fluorescence and photoacoustic properties of our DNA nanostructures as a function of fluorophore-quencher separation distance. Importantly, we demonstrate experimentally the potential of PAT for performing nanoscale distance measurements at depth in tissue mimicking phantoms and also highlight the utility of DNA nanostructures to enhance the photoacoustic signal generation capabilities of small molecule fluorescent dyes.

## RESULTS AND DISCUSSION

In our experimental realization, we used DNA nanostructures that consist of double-stranded single helices carrying an NIR fluorophore (either IRDye 800CW or Cy5.5) and quencher (IRDye QC-1) pair at six different distances (see Figure S1 for the dye and quencher chemical structures). The distance between the dye and quencher was controlled by varying the number (**N**) of nucleotides (**nts**) between them (series **Nnts-DQ**, illustrated in Figure 1). We also prepared two additional series as controls (see Supporting Information section S1), carrying either the fluorophore alone (series **Nnts-D**) or the quencher alone (series **Nnts-Q**). The sequences, layout and assignment of all oligonucleotides composing the DNA nanostructures are shown in Figure S2 and Table S1. The nanostructures were prepared in phosphate buffered saline (PBS) and were analyzed with gel electrophoresis to confirm their correct folding (Figure S3).

**Figure 1.**
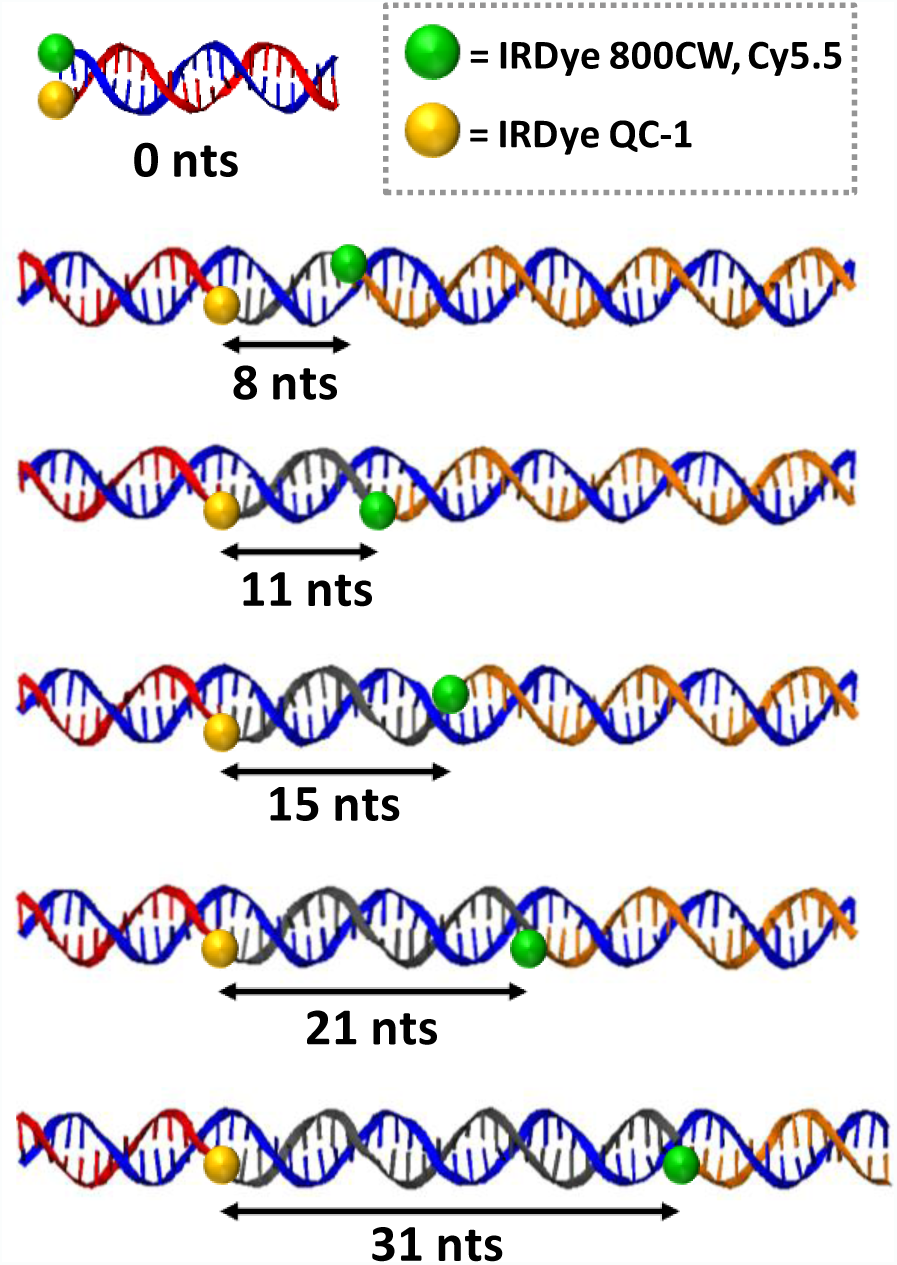
Schematics of the reported DNA nanostructures. Each strand contained in each of the nanostructures is represented with a different color. IRDye QC-1 quencher is represented as yellow spheres. Fluorophores (IRDye 800CW or Cy5.5) are shown as a green spheres. The number of nucleotides separating the quencher and the dye on each design is represented with an arrow. The dye and the quencher are connected at the terminal part of the oligonucleotides. Series Nnts-DQ possess both the dye and quencher, series Nnts-D only dye and series Nnts-Q only the quencher. Nts = Nucleotides.

Optical characterization of the nanostructures with N= 8 to 31nts confirmed their absorbance and distance-dependent fluorescence quenching behaviors. All characterization measurements were performed using 2 μM DNA concentration for each of the nanostructures prepared separately and were averaged over 3 replicates. The absorbance measurements for **Nnts-DQ** containing IRDye 800CW (Figure 2a) and Cy5.5 (Figure 2b) show no changes in the peak absorbance wavelengths for **Nnts-DQ** (solid-dark line) when compared to **Nnts-D** (solid-light line) and **Nnts-Q** (dotted line) nanostructures.

**Figure 2.**
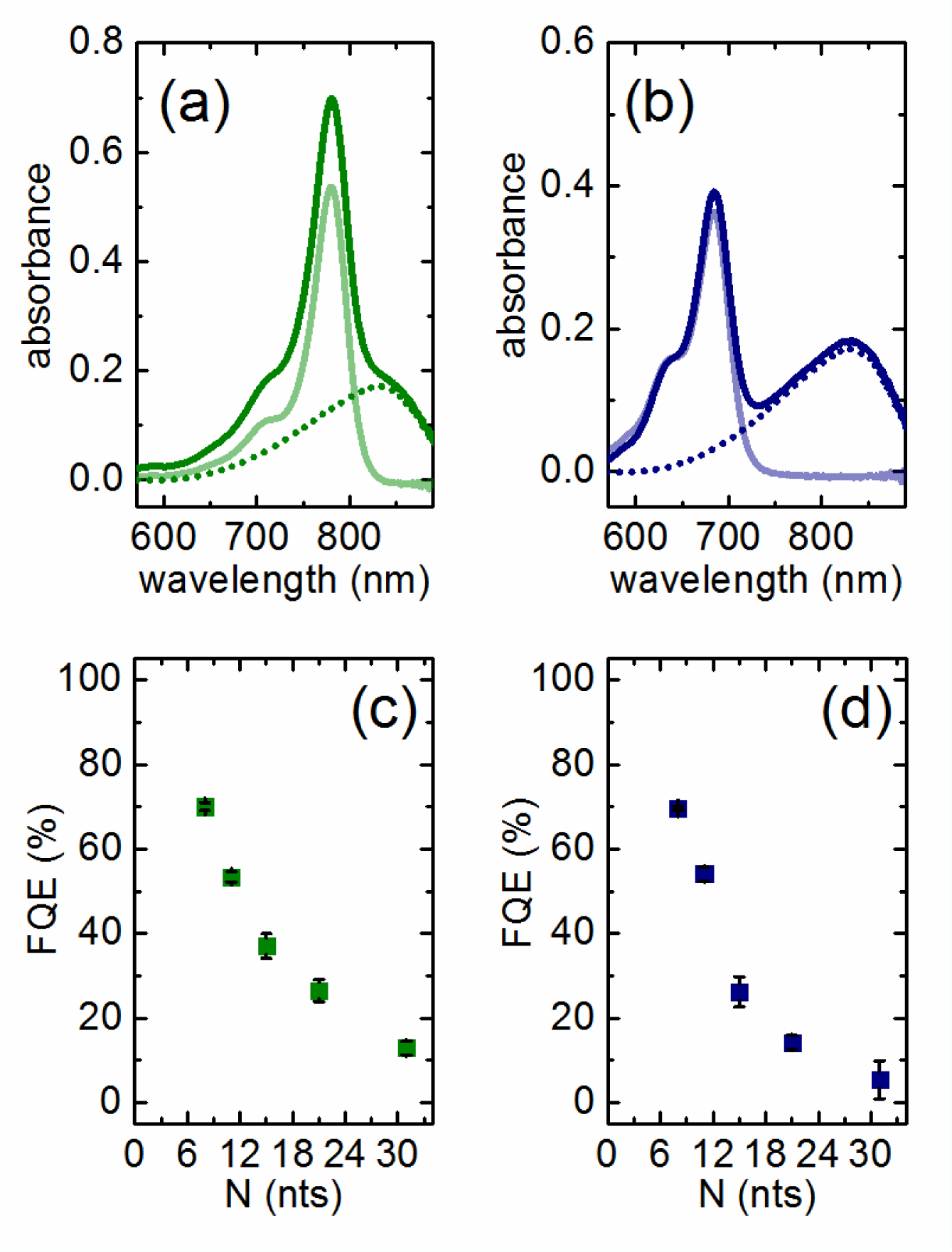
Optical characterization of the DNA nanostructures with spacing of N= 8, 11, 15, 21 and 31 nts. (a) and (b) Absorbance spectra of the series Nnts-DQ (solid-dark line), Nnts-D (solid-light line) and Nnts-Q (dotted line) for N=15nts derivatives (as example of the series) of (a) IRDye 800CW nanostructures and (b) Cy5.5 nanostructures. (c) and (d) Distance-dependent quenching efficiency of emission obtained as a function of the number of nucleotides (N) between the quencher and the dye in DNA designs for (c) IRDye 800CW derivatives and (d) Cy5.5 derivatives. Quenching efficiency is calculated as described in the text. Each data point in (c) and (d) corresponds to the average of the values obtained from 3 replicates. The error bar is the propagation of error obtained from the calculations performed to determine the FQE.

Fluorescence emissions from each of the nanostructures were measured at their corresponding peak absorbance wavelengths (Table S2) to evaluate distance-dependent fluorescence quenching behavior. For the five different **Nnts-DQ** nanostructures, this behavior is described in terms of the fluorescence quenching efficiency (**FQE**) which is given as **(FQE= 100 × [(I_D_- I_DQ_)/(I_D_)])**, where **I_D_** and **I_DQ_** are the peak emission intensities obtained from the **Nnts-D** and **Nnts-DQ** nanostructures respectively. Emission measurements obtained from **Nnts-DQ** nanostructures clearly show enhanced FQE associated with corresponding shortening of the distance between the fluorophore and quencher (IRDye 800CW, Figure 2c; Cy5.5 Figure 2d). These relationships show r^2^ values of 0.987 (IRDye 800CW) and 0.997 (Cy5.5) upon fitting to the Förster Resonance Energy Transfer (FRET) equation (Supporting Information section S2, Figure S4), indicating that the physical origin of the quenching mechanism in these nanostructures is likely to be due to FRET.

Interestingly, optical characterization of the **Nnts-DQ** nanostructures with the shortest separation (N= 0nts) showed a markedly different optical response, inconsistent with a quenching behaviour based on FRET. The absorbance spectrum obtained from the **0nts-DQ** nanostructures carrying IRDye 800CW (full dark green line, Figure 3a) showed an additional peak at 719 nm. An equivalent blue-shifted absorbance peak (at 664 nm) was also observed in the case of the **0nts-DQ** with Cy5.5 (Figure 3b). The fluorescence emission measurements obtained from these nanostructures indicate that the FQE for the **0nts-DQ** nanostructures is extremely high (above 95%) for both classes of fluorophore-quencher pairs (IRDye 800CW, Cy5.5; Figure 3e). The observed modification of the absorbance spectra, together with the high FQE value, suggest a static quenching mechanism for **0nts-DQ** nanostructures produced by the stacking of the fluorophore and quencher.^33^

**Figure 3.**
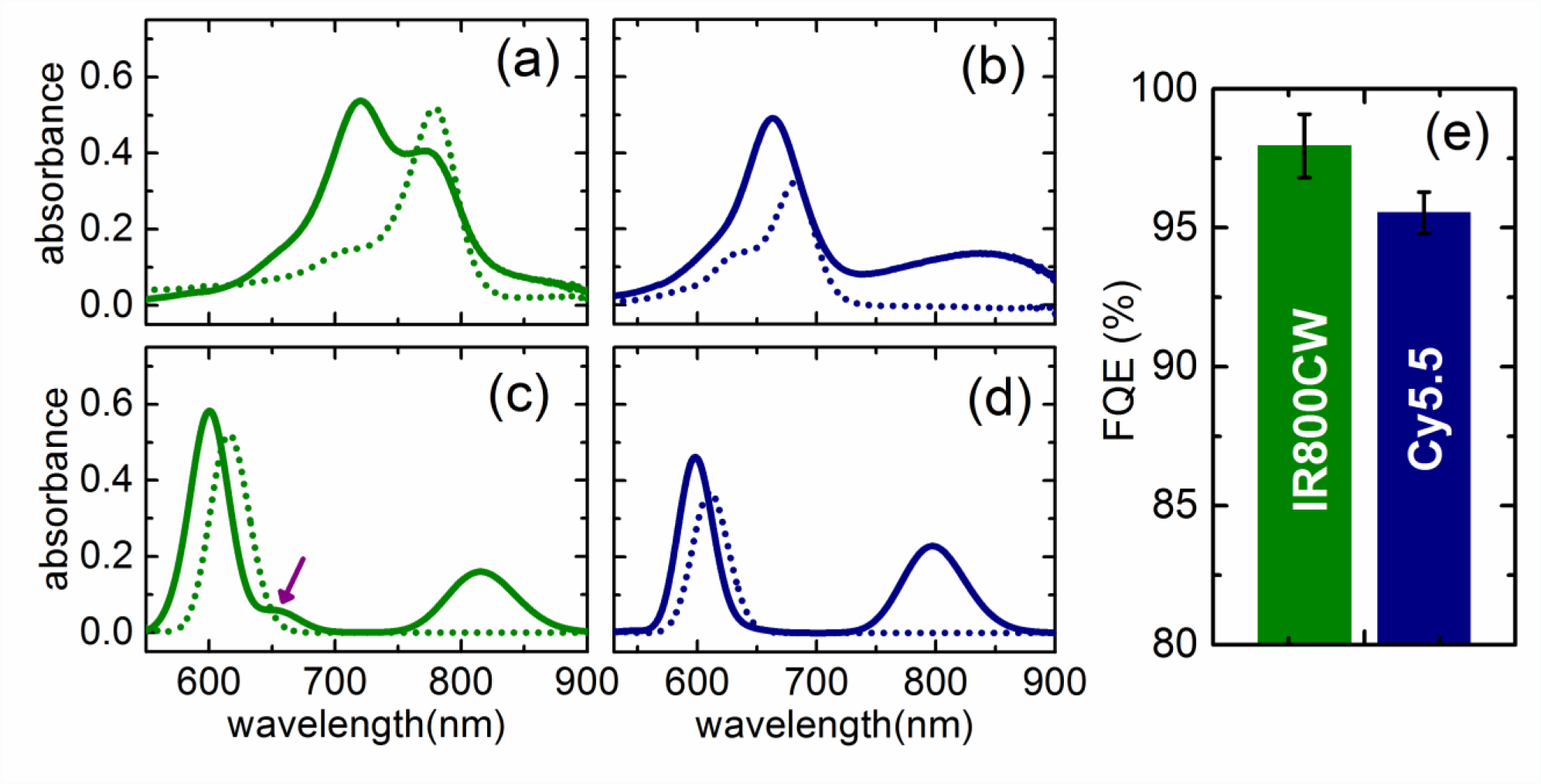
Optical characterization of the 0 nts DNA nanostructures. (a) and (b) Experimental absorbance spectra. (c) and (d) Theoretical absorbance spectra as predicted by TDDFT calculations. IR800CW and Cy5.5 derivatives are shown in green and blue respectively. 0nts-DQ (solid line), 0nts-D (dotted line). (e) Fluorescence quenching efficiency obtained from the 0nts DNA designs. Quenching efficiency is calculated as described in the text. FQE data corresponds to the average of the values obtained in three individual samples. The error bar is the propagation of error obtained from the calculations performed to determine the FQE.

In order to confirm the origin of the changes in absorbance spectra for **0nts-DQ** nanostructures, we built *in silico* model structures for **0nts-D**, **0nts-Q** and stacked fluorophore-quencher systems (**0nts-DQ**) for both IRDye 800CW/IRDye QC-1 and Cy5.5/IRDye QC-1 and computed the corresponding absorbance spectra using time-dependent density-functional theory (TDDFT) (see Materials and Methods for information on the computational details and the construction of stacked fluorophore-quencher models).^34^ The resultant predicted spectra in comparison with the experimental results for N= 0 nts are shown in Figure 3c and d and Figure S5. While the theoretical results overestimate the blue-shift of the main absorption peak of the fluorophore with respect to the quencher for both IRDye 800CW and Cy5.5, the spectral anomalies of **0nts-DQ** with respect to larger separations are correctly reproduced by the stacked fluorophore-quencher models. Most notably, the TDDFT results correctly predict a blue-shift of the main absorption peak of the fluorophore, as well as a significant drop in the peak absorbance associated with IRDye QC-1. Furthermore, the theoretical results for **0nts-DQ** with IRDye 800CW also show a red-shifted second peak next to the absorbance maximum (∼650 nm, see purple arrow in Figure 3c), although the peak height is lower than in the experimental results. The TDDFT results provide strong evidence to support the hypothesis that the spectral changes of **0nts-DQ** with respect to those obtained for larger separations are indeed due to a dipole coupling of the dominant excited states of the fluorophore and the quencher facilitated by a stacked conformation.^35^

We next investigated the effect of fluorescence quenching on PA signal generation under tissue mimicking conditions. The DNA nanostructures were encapsulated within thin-walled plastic straws at 1 cm depth in tissue mimicking phantoms and photoacoustic imaging data was obtained using a commercial PAT system (see Materials and Methods). Photoacoustic signals were acquired from 3 replicates of separately prepared nanostructures, using multiple excitation wavelengths and scan positions for each phantom. Quantification of the photoacoustic signals was performed by extracting the mean pixel intensity (MPI) from a region of interest (ROI) drawn within the straw position in the reconstructed images at different wavelengths (see Figure S6 and Table S3). Photoacoustic signal enhancement (PE) was then quantified as **PE= 100 × {[I_DQ_- (I_D_+I_Q_)]/ (I_D_+I_Q_)}**, where **I_DQ_**, **I_D_** and **I_Q_** are the averaged MPIs measured from the nanostructures of series **Nnts-DQ, Nnts-D** and **Nnts-Q** respectively. PE values were calculated from the data extracted at the scanned wavelengths at which the absorbance for the Nnts-DQ nanostructures were maximum (Table S4).

The PE obtained from **Nnts-DQ** nanostructures containing IRDye 800CW (Figure 4a) and Cy5.5 (Figure 4b) show that significant enhancements in photoacoustic signal are indeed observed in a distance-dependent manner (see also Figure S6 and S7). The results therefore show a direct dependence of photoacoustic signal on fluorescence quenching, where an increased level of fluorescence quenching directly contributes to a substantial enhancement in photoacoustic signals. Of particular note are the extremely high levels of PE obtained from **0nts-DQ** nanostructures (Figure 4c) which had the shortest molecular separation and the highest fluorescence FQE (Figure 3e). The highest level of PE occurs in the blue shifted peak identified by absorbance measurements (Figure 3a,b and Figure S6). These results indicate that PAT could be used to monitor distance-dependent interactions at depth in living subjects.

**Figure 4.**
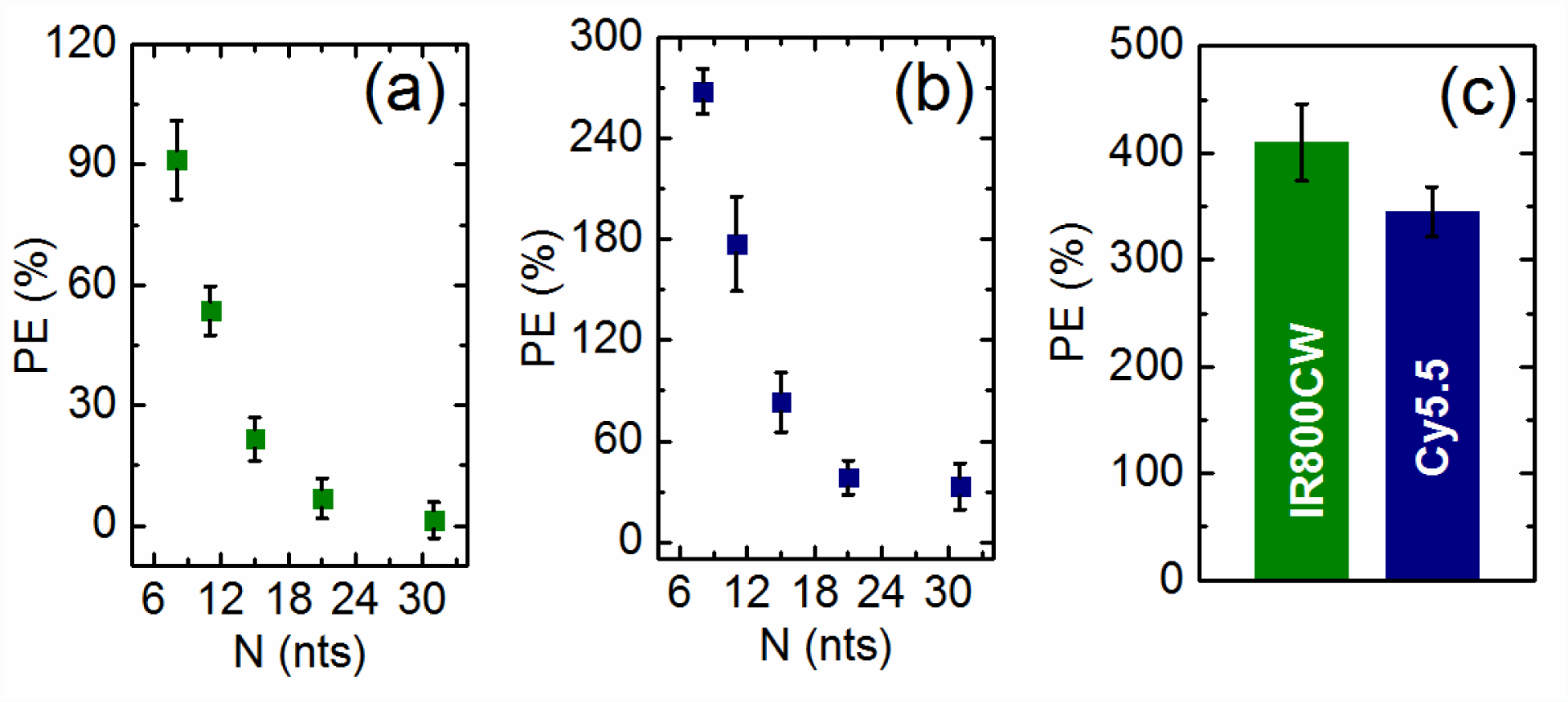
Photoacoustic response of the DNA nanostructures. (a) and (b) PE is given as a function of the number of nucleotides (N) between dye and quencher for designs with N= 8, 11, 15, 21 and 31 nts for (a) IRDye 800CW derivatives and (b) Cy5.5 derivatives. (c) PE obtained from the 0 nts nanostructures. Each data point in (a)-(c) corresponds to the average of the values obtained from 3 replicates. The error bar is the propagation of error obtained from the calculations performed to determine the PE. The reported PE values for the nanostructures with N= 8, 11, 15, 21 and 31 nts were calculated at wavelengths of 778 nm and 682 nm for IRDye 800CW and Cy5.5 derivatives respectively. PE values for nanostructures with N= 0 nts were calculated at 719 nm and 665 nm for IRDye 800CW and Cy5.5 derivatives respectively (see Table S4 in the Supporting Information).

## CONCLUSIONS

To summarize, we have shown that photoacoustic signal enhancements can be precisely tuned by controlling the distance between a fluorophore and a quencher. The mechanism of this process is related to fluorescence quenching; it occurs primarily via a FRET mechanism for nanostructures N=8 to 31nts, but is more likely associated with static quenching at N= 0 nts due to stacking of the fluorophore and quencher molecules. The direct demonstration of the link between photoacoustic signal generation and the non-radiative decay of absorbed optical energy due to FRET establishes a new strategy to probe molecular scale distance-dependent measurements at centimeter depths, which could be readily extended into deep tissue imaging *in vivo*. In addition, the high photoacoustic signal enhancement provided by the N= 0 nts nanostructure could be exploited to create a biodegradable contrast agent based on small molecule fluorescent dyes. Targeting this nanostructure to a specific biochemical process would enable highly efficient photoacoustic molecular imaging with translational potential.^32^ In conclusion, we have shown that the process of fluorescence quenching can be exploited to create photoacoustic rulers, which could, in the future, be applied for studies of molecular interactions at depth in living subjects.

## MATERIALS AND METHODS

### DNA nanostructures design, assembly and characterization

The oligonucleotides were purchased from IDT (Integrated DNA Technologies, Inc). The sequences were randomly generated and NUPACK^36^ was used to check that they were appropriate to minimize the formation of homodimers or hairpins. Complementary strands were obtained using the open source DNA origami software caDNAno.^37^ The DNA nanostructures were assembled in phosphate buffered saline (pH=7.4) to a final concentration of 2μM DNA. The mixture was subjected to thermal-annealing for 45 min to ensure maximum yield in the folding. The heating program utilized was from 70 to 25°C in 90 steps (0.5°C per step, 30s each step) and finally samples were kept at 4°C. Polyacrylamide gel electrophoresis (PAGE) was performed to confirm the correct assembly. PAGE (10%) was prepared and run in a solution containing 11mM MgCl_2_ and buffered with 0.5xTBE (pH = 8.3). 50bp DNA Ladder (Invitrogen, Thermo Fisher Scientific Inc.) was used as reference. The samples were run at 100 V for 90min. The gels were stained with GelRed and visualized using a UVP gel doc-it imaging system (Figure S3).

### Absorbance and emission measurements

The absorbance and fluorescence emission of the DNA nanostructures were measured at 34°C (to mimic PAT measurements temperature conditions) with a fixed concentration of 2μM of DNA. Absorbance and emission properties were measured using a UV-Vis Spectrophotometer (Varian Cary 300 Bio, Agilent Technologies, Inc.) and Fluorescence Spectrophotometer (Varian Cary Eclipse, Agilent Technologies, Inc.) respectively. Absorbance and emission data were measured at 1 nm steps. Fluorescence excitation was performed at the wavelengths detailed in Table S2, which corresponds to the respective absorbance maxima. Note that different excitation wavelengths were used for N= 8 to 31 nts and N=0 nts nanostructures due to the different maxima observed in the absorbance spectra.

**Time-dependent density functional theory (TDDFT) calculations**

TDDFT calculations were performed on reduced models of IRDye QC-1, IRDye 800CW and Cy5.5, while the effects of the DNA backbone were ignored (Figure S8). The initial structures of the isolated dyes were prepared using the BOSS software^38^ and then reoptimized using DFT at the PBE^39^ level of theory. Absorption spectra of IRDye QC-1, IRDye 800CW and Cy5.5 in isolation were calculated using an implicit solvation model with a relative dielectric constant of 80 in order to account for the screening of the aqueous environment.^40^

The two models of the stacked dye-quencher systems for a nucleotide separation of zero were obtained by taking the optimized isolated structures of IRDye QC-1, as well as IRDye 800CW and Cy5.5, and placing the dye on top of the quencher in a flat stacking, such that the alignment of the dipole moments of the dominant excitations in the individual systems is maximized. It was found that the closest stacking expected to maximize the excitonic coupling between the dye and the quencher and thus the largest changes in the absorption spectra can be achieved by rotating the quencher by 180 degrees with respect to the dye system. The initial structures for IRDye 800CW/ IRDye QC-1 and Cy5.5/IRDye QC-1 were optimized using DFT at the PBE level, where the van-der-Waals interactions between the dye and the quencher were accounted for by the empirical dispersion correction of Wu and Yang^41^ (see Figure S9 for final structures of the resulting combined systems). TDDFT calculations were then performed on the two optimized, stacked dye-quencher systems, again using an implicit solvation model to account for the dielectric screening of the aqueous environment.

All DFT and TDDFT calculations were performed using the ONETEP code.^42-44^ A 800 eV kinetic energy cutoff on the underlying psinc basis set and a 10 a_0_ cutoff radius on all localized support functions was used throughout. All calculations were performed using the PBE functional and norm-conserving pseudopotentials. The calculation settings chosen in this work have been previously shown to yield fully converged excitation energies for small to medium sized chromophores in vacuum and solution.^34, 44^

### Photoacoustic tomography (PAT) measurements and calculations

Photacoustic measurements were performed using a commercial PAT system (inVision256-TF; iThera Medical GmbH) and tissue mimicking phantoms that closely mimic the optical and acoustic properties of biological tissues (see schematic representation in Figure S10). The commercial PAT system that has been described previously^45, 46^ uses a tunable (660–1300nm) optical parametric oscillator pumped by a nanosecond pulsed Nd:YAG laser to provide 9ns excitation pulses at 10Hz repetition rate. Ten arms of a fiber bundle illuminate a ring of ∼8mm width around the sample. The phantom was mounted in a motorized holder for linear translation in the z-direction over a range of <150mm. Acoustic coupling between the phantom and ultrasound transducers was achieved using a temperature maintained imaging chamber, filled with degassed, deionized water. For ultrasound detection, 256 toroidally focused ultrasound transducers specified at 5MHz center frequency, 60% bandwidth, are organized in a concave array with 270 degree angular coverage and a radius of curvature of 4 cm.

The phantoms were fabricated using agar as the base material; nigrosin dye and intralipid were added to provide an absorption coefficient of 0.05cm^-1^ and reduced scattering coefficient of 5cm^-1^ according to our standard procedure.^47^ All the reagents for phantoms fabrication were purchased from Sigma Aldrich (Sigma-Aldrich Co.) unless otherwise stated. The DNA nanostructures were suspended inside sealed thin walled plastic tubes (0.3 cm diameter) that were placed at the center of the cylindrical phantoms (2 cm diameter) as shown in Figure S10b. We have shown previously that these conditions accurately mimic the optical properties of mouse tissue.^47^ For all the measurements, the phantoms were maintained at 34°C inside the water bath. PAT data were acquired at the specific excitation wavelengths (see Table S3) with 10 time frames averaging and at 5 scan locations separated by a 1 mm step size for averaging over position. A model-based reconstruction algorithm^48^ was used to reconstruct the PAT images and PA data were extracted at different wavelengths as shown in Figure S6. Mean pixel intensity (MPI) values were extracted from a region of interest (ROI) drawn within the thin walled plastic straw and the averaged values over the 5 scan positions were used for further analysis.

## ACKNOWLEDGMENT

JJ, JW and SEB are funded by the EPSRC-CRUK Cancer Imaging Centre in Cambridge and Manchester (C197/A16465); CRUK (C14303/A17197, C47594/A16267) and the EU-FP7- agreement FP7-PEOPLE-2013-CIG-630729. K. N. B. acknowledges the ERASMUS placement organization for ERASMUS+ funding. P. K. acknowledges funding by the EPSRC. T. J. Z. acknowledges the support of EPSRC Grant EP/J017639/1. S.H.A. acknowledges support from a Herchel Smith postdoctoral fellowship and from a Cancer Research UK Cambridge Centre Research Grant. The authors thank Professor Ulrich Keyser for valuable inputs.

## REFERENCES

1. Jares-Erijman, E. A.; Jovin, T. M. Nat. Biotech. 2003, 21, (11), 1387–1395.

2. Anker, J. N.; Hall, W. P.; Lyandres, O.; Shah, N. C.; Zhao, J.; Van Duyne, R. P. Nat. Mat. 2008, 7, (6), 442–453.

3. Taruttis, A.; Ntziachristos, V. Nat. Photonics 2015, 9, (4), 219–227.

4. Piston, D. W.; Kremers, G.-J. Trends Biochem. Sci. 2007, 32, (9), 407–414.

5. Gaits, F.; Hahn, K. Sci. STKE 2003, 2003, (165), pe3–pe3.

6. Hirata, E.; Girotti, M. R.; Viros, A.; Hooper, S.; Spencer-Dene, B.; Matsuda, M.; Larkin, J.; Marais, R.; Sahai, E. Cancer Cell 2015, 27, (4), 574–588.

7. Förster, T. Annalen der physik 1948, 437, (1-2), 55–75.

8. Aoki, K.; Kamioka, Y.; Matsuda, M. Dev Growth Differ. 2013, 55, (4), 515–522.

9. Mächler, P.; Wyss, M. T.; Elsayed, M.; Stobart, J.; Gutierrez, R.; von Faber-Castell, A.; Kaelin, V.; Zuend, M.; San Martín, A.; Romero-Gómez, I. Cell metabolism 2016, 23, (1), 94–102.

10. Bouchaala, R.; Mercier, L.; Andreiuk, B.; Mély, Y.; Vandamme, T.; Anton, N.; Goetz, J. G.; Klymchenko, A. S. J. Controlled Release 2016, 236, 57–67.

11. Gravier, J.; Sancey, L.; Hirsjärvi, S.; Rustique, E.; Passirani, C.; Benoît, J.-P.; Coll, J.-L.; Texier, I. Molecular pharmaceutics 2014, 11, (9), 3133–3144.

12. Wang, L. V.; Hu, S. Science 2012, 335, (6075), 1458–1462.

13. Wang, L. V. IEEE Journal of Selected Topics in Quantum Electronics 2008, 14, (1), 171–179.

14. Chance, R.; Prock, A.; Silbey, R. Adv. Chem. Phys 1978, 37, (1), 65.

15. Pinheiro, A. V.; Han, D.; Shih, W. M.; Yan, H. Nat. Nanotech. 2011, 6, (12), 763–772.

16. Bandy, T. J.; Brewer, A.; Burns, J. R.; Marth, G.; Nguyen, T.; Stulz, E. Chem. Soc. Rev. 2011, 40, (1), 138–148.

17. Acuna, G. P.; Bucher, M.; Stein, I. H.; Steinhauer, C.; Kuzyk, A.; Holzmeister, P.; Schreiber, R.; Moroz, A.; Stefani, F. D.; Liedl, T. ACS Nano 2012, 6, (4), 3189–3195.

18. Voigt, N. V.; Tørring, T.; Rotaru, A.; Jacobsen, M. F.; Ravnsbæk, J. B.; Subramani, R.; Mamdouh, W.; Kjems, J.; Mokhir, A.; Besenbacher, F. Nat. Nanotech. 2010, 5, (3), 200–203.

19. Massey, M.; Algar, W. R.; Krull, U. J. Anal. Chim. Acta 2006, 568, (1), 181–189.

20. Hemmig, E. A.; Creatore, C.; Wünsch, B.; Hecker, L.; Mair, P.; Parker, M. A.; Emmott, S.; Tinnefeld, P.; Keyser, U. F.; Chin, A. W. Nano Lett. 2016, 16, (4), 2369–2374.

21. Kato, T.; Kashida, H.; Kishida, H.; Yada, H.; Okamoto, H.; Asanuma, H. J. Am. Chem. Soc. 2013, 135, (2), 741–750.

22. Sindbert, S.; Kalinin, S.; Nguyen, H.; Kienzler, A.; Clima, L.; Bannwarth, W.; Appel, B.; Müller, S.; Seidel, C. A. J. Am. Chem. Soc. 2011, 133, (8), 2463–2480.

23. Li, C.-Y.; Hemmig, E. A.; Kong, J.; Yoo, J.; Hernández-Ainsa, S.; Keyser, U. F.; Aksimentiev, A. ACS Nano 2015, 9, (2), 1420–1433.

24. Didenko, V. V. Biotechniques 2001, 31, (5), 1106.

25. Stein, I. H.; Schüller, V.; Böhm, P.; Tinnefeld, P.; Liedl, T. ChemPhysChem 2011, 12, (3), 689–695.

26. Tsukanov, R.; Tomov, T. E.; Liber, M.; Berger, Y.; Nir, E. Acc. Chem. Res. 2014, 47, (6), 1789–1798.

27. White, S. S.; Balasubramanian, S.; Klenerman, D.; Ying, L. Angew. Chem. Int. Ed. 2006, 45, (45), 7540–7543.

28. Gietl, A.; Holzmeister, P.; Grohmann, D.; Tinnefeld, P. Nucleic Acids Res. 2012, 40, (14), e110–e110.

29. Nickels, P. C.; Wünsch, B.; Holzmeister, P.; Bae, W.; Kneer, L. M.; Grohmann, D.; Tinnefeld, P.; Liedl, T. Science 2016, 354, (6310), 305–307.

30. Andersen, E. S.; Dong, M.; Nielsen, M. M.; Jahn, K.; Subramani, R.; Mamdouh, W.; Golas, M. M.; Sander, B.; Stark, H.; Oliveira, C. L.; Pedersen, J. S.; Birkedal, V.; Besenbacher, F.; Gothelf, K. V.; Kjems, J. Nature 2009, 459, (7243), 73–76.

31. Tomov, T. E.; Tsukanov, R.; Liber, M.; Masoud, R.; Plavner, N.; Nir, E. J. Am. Chem. Soc. 2013, 135, (32), 11935–11941.

32. Weber, J.; Beard, P. C.; Bohndiek, S. E. Nat. Methods 2016, 13, (8), 639–650.

33. Marras, S. A. Fluorescent Energy Transfer Nucleic Acid Probes: Designs and Protocols 2006, 3–16.

34. Zuehlsdorff, T. J.; Haynes, P. D.; Hanke, F.; Payne, M. C.; Hine, N. D. J. Chem. Theory Comput. 2016, 12, (4), 1853–1861.

35. Nicoli, F.; Roos, M. K.; Hemmig, E. A.; Di Antonio, M.; de Vivie-Riedle, R.; Liedl, T. J. Phys. Chem. A 2016, (50), 9941–9947.

36. Zadeh, J. N.; Steenberg, C. D.; Bois, J. S.; Wolfe, B. R.; Pierce, M. B.; Khan, A. R.; Dirks, R. M.; Pierce, N. A. J. Comput. Chem. 2011, 32, (1), 170–173.

37. Douglas, S. M.; Marblestone, A. H.; Teerapittayanon, S.; Vazquez, A.; Church, G. M.; Shih, W. M. Nucleic Acids Res. 2009, 37, (15), 5001–5006.

38. Jorgensen, W. L.; Tirado–Rives, J. J. Comp. Chem. 2005, 26, (16), 1689–1700.

39. Perdew, J. P.; Burke, K.; Ernzerhof, M. Phys. Rev. Lett. 1996, 77, (18), 3865.

40. Dziedzic, J.; Helal, H. H.; Skylaris, C.-K.; Mostofi, A. A.; Payne, M. C. EPL (Europhysics Letters) 2011, 95, (4), 43001.

41. Wu, Q.; Yang, W. J. Chem. Phys. 2002, 116, (2), 515–524.

42. Zuehlsdorff, T. J.; Hine, N. D.; Spencer, J. S.; Harrison, N. M.; Riley, D. J.; Haynes, P. D. J. Chem. Phys. 2013, 139, (6), 064104.

43. Skylaris, C.-K.; Haynes, P. D.; Mostofi, A. A.; Payne, M. C. J. Chem. Phys. 2005, 122, (8), 084119.

44. Zuehlsdorff, T.; Hine, N.; Payne, M. C.; Haynes, P. D. J. Chem. Phys. 2015, 143, (20), 204107.

45. Morscher, S.; Driessen, W. H.; Claussen, J.; Burton, N. C. Photoacoustics 2014, 2, (3), 103–110.

46. Dima, A.; Burton, N. C.; Ntziachristos, V. J. Biomed. Opt. 2014, 19, (3), 036021–036021.

47. Joseph, J.; Tomaszewski, M.; Quiros-Gonzalez, I.; Weber, J.; Brunker, J.; Bohndiek, S. E. J. Nuc. Med. 2017, jnumed. 116.182311.

48. Dean-Ben, X. L.; Buehler, A.; Ntziachristos, V.; Razansky, D. IEEE Transactions on Medical Imaging 2012, 31, (10), 1922–1928.

